# External and Internal Load Profiles of Male and Female Participants during a Walking Football Practice

**DOI:** 10.1101/2025.01.15.633234

**Authors:** Júlio A. Costa, Catarina Pereira, Ana Barbosa, André Seabra, João Brito, Ana Pinto, Catarina Martins, Rafaela Moreira, Bruno Gonçalves

**Affiliations:** Portugal Football School, Portuguese Football Federation, FPF, Oeiras, Portugal; Departamento de Desporto e Saúde, Escola de Saúde e Desenvolvimento Humano, Universidade de Évora, Évora, Portugal; Comprehensive Health Research Centre (CHRC), Universidade de Évora, Évora, Portugal; EPIUnit ITR, Institute of Public Health of the University Porto, University of Porto, Rua das Taipas, n° 135, 4050-600 Porto, Portugal

**Keywords:** Aging, cardiovascular health, gender, physical fitness, exercise

## Abstract

**BACKGROUND:** High levels of sedentary behavior among older adults highlight the importance of walking football (WF) in promoting physical activity for healthy aging. This study examines the external and internal load profiles of male and female participants during a WF tournament, addressing a gap in research on game demands and induced load.

**METHODS:** The study involved 176 players aged 50+ participating in a 40-min, 5v5 WF tournament with unlimited substitutions. External load (total and categorized distances) was measured using Global Positioning System (GPS), while heart rate (HR) monitors assessed internal load, including absolute HR and intensity zones based on %HRmax.

**RESULTS:** The proportion of male participants (n=123; 70.3%) was higher than females (n=52; 29.7%), p<.001. They were similar in age (61.6±8.6 and 60.8±6.9, respectively). Males covered a higher distance per minute than females, with sex showing a moderate effect (63.3±10.7 m/min vs. 54.7±15.8 m/min; p < .001; Cohen’s *d*_*unbiased*_ = 0.69 [0.36; 1.03]), especially in fast walking (41.7±12.2 m/min vs. 32.6±16.7 m/min; p < .001; Cohen’s *d*_*unbiased*_ = 0.66 [0.33; 1.00]). Males played more time than females (22:26±09:47 min:ss vs. 15:41±07:46 min:ss; p<.001), with moderate effect (Cohen’s *d*_*unbiased*_ = 0.73 [0.40; 1.06]). However, no differences between sexes were identified in the internal load variables, such that the female average %HRmax was 80±11% and the male was 82±8% during the practice.

**CONCLUSIONS:** Overall, while males generally exhibit higher external loads in WF, both sexes experience similar internal load demands, highlighting WF’s potential as a scalable, health-promoting intervention for aging populations.

## Introduction

Literature reports a substantial growth in the body of evidence on the multiple health and well-being benefits of different types, intensities, and durations of physical activity, as well as the negative impact of sedentary behavior on health [1]. Physical activity in older ages is a known protective factor for mental health and prevention and management of noncommunicable diseases, such as cardiovascular disease, type 2 diabetes, or cancer [1, 2] Physical activity also improves functional fitness, mobility and ability, therefore contributing to older adults independence and well-being [3].

Several international organizations, such as the World Health Organization (WHO) [1], the American College of Sports Medicine (ACSM) [4], and the Centers for Disease Control and Prevention (CDC) [5], recommend that adults engage in at least 150 minutes of moderate-intensity physical activity or 75 minutes of vigorous-intensity physical activity per week, or an equivalent combination of moderate- and vigorous-intensity aerobic activity, spread throughout the week. Older adults should follow the guidelines for adults and add multicomponent physical activity that includes flexibility and balance exercises on three or more days a week, to enhance functional capacity and prevent falls [6].

Walking, depending on its intensity and duration, can play a significant role in promoting health among older adults [2]. Although some authors report a modest improvement over time in older adults’ adherence to physical activity recommendations, they also note that adherence remains low [7], with older adults exhibiting high levels of physical inactivity that tend to increase with age [8]. Given this scenario, the WHO established a global action plan on physical activity for 2018-2030, in which action 3.4 Intent to “enhance the provision of and opportunities for, appropriately tailored programs and services aimed at increasing physical activity and reducing sedentary behavior…” [9].

Walking Football (WF), a modified format of soccer, has been growing among individuals aged 50 years and older, emphasizing inclusivity and safety by prohibiting running and physical contact [10]. Its development has been driven by the increasing recognition of physical activity as a critical determinant of health, particularly for aging populations [11]. WF has emerged as an innovative solution, offering older adults a light-to-vigorous intensity activity with reduced injury risks [12]. Existing literature supports its potential for improving physical fitness and promoting social connectedness among older adults [11, 13, 14]. However, the physiological demands and outcomes of WF remain underexplored, particularly in terms of sex-based differences in internal and external load metrics. This study addresses these gaps, particularly the physiological profiles and WF practice demands of male and female WF participants in a WF tournament context.

External load metrics, such as distance covered and movement speeds, offer insights into the physical demands of the modality [11]. Regarding internal load measures, including heart rate (HR) zones, provide a window into cardiovascular responses [13, 14]. While studies have shown that WF can elicit health-enhancing physical activity levels [11, 13, 14], there remains a scarcity of data comparing these responses between male and female participants.

Therefore, this study investigates sex-specific differences in WF by analyzing external and internal load profiles during a friendly tournament. It was hypothesized that males would exhibit higher external loads (e.g., distance and speed) than females, while internal loads (%HRmax) would remain similar due to the consistent moderate-intensity demands of WF.

## Methods

### Study Design

This study utilized a cross-sectional observational analytical design to assess participants’ external and internal load profiles in WF games during a national WF friendly tournament held in Oeiras, Portugal, in 2023.

### Participants

Participants were eligible for the study if they were aged ≥ 50 years and provided a valid medical fitness certificate confirming their suitability for sports participation. The recruitment process, conducted by the project manager via email and in person, took place between June 1^st^ and July 31^st^ 2023 and targeted each football association in Portugal.

Participants provided written informed consent prior to participation. Ethical approval was obtained from the Portugal Football School Ethics Committee (Protocol number: 21/CEPFS/2023), and the study was conducted in compliance with the Declaration of Helsinki.

### Procedures

WF games were played under standardized rules tailored for participants over 50 years of age: no running with or without the ball; maximum of three touches on the ball by player, no physical contact, including slide tackles [10]. Besides these rules, it was defined that the ball must always be played below the players’ average waist height [12]. Games were held on a 40×20 meters and lasted 40 minutes, divided into two 20-minute halves with a 10-minute interval. Teams consisted of five outfield players, and substitutions were unlimited. Each team played four games.

Games were played in an outdoor natural grass. Data were collected across all 12 games of the tournament by trained researchers with a Human Kinetic Sciences background. Each researcher performed always the same evaluation task.

### External load

External load metrics were captured using 10Hz Global Positioning System (GPS) devices (STATSports Apex, Northern Ireland). The devices were turned on 10 min before the WF games and placed on the participants who wore a customized and specific neoprene vest in the midline between the scapulae at the level of the seventh cervical vertebra (C7). Distances were categorized by speed zones distance covered as: distance covered at speeds < 4 km/h (low-intensity activity); distance covered at speeds > 4 km/h (higher-intensity activity)[11, 15]. The reliability and validity of the GPS devices were previously established in sports performance studies [16].

### Internal load

To measure internal load during WF games, a GARMIN HR (Garmin Ltd., Olathe, Kansas, United States) monitoring band was used by the participants [17]. Absolute HR values were recorded and the percentage of age-estimated maximal HR (%HRmax) was calculated using the equation HRmax=211−0.64×age [18]. However, if players reached HR values during the game that exceeded this predicted maximum, those observed values were instead used as their HRmax. Average HR, % HRmax, HRpeak were registered, and the player’s HR data were categorized into intensity zones: Zone 1: <50% HRmax; Zone 2: 50–60% HRmax; Zone 3: 60– 70% HRmax; Zone 4: 70–80% HRmax; Zone 5: 80–90% HRmax; Zone 6: >90% HRmax [11] (adapted for walking football older participants)[15].

### Statistical Analysis

After preliminary inspections for distribution and assumptions, an independent sample t-test analysis was processed to identify the effect of the sex (female vs male) on the considered variables. The statistical analysis was performed using the Statistical Package for the Social Sciences software (SPSS, Inc., Chicago, IL, USA), and statistical significance was set at p < .05.

An estimation techniques approach was carried to overcome the shortcomings associated with traditional N-P null hypothesis significance testing [19, 20]. The Cohen’s *d*_*unbiased*_ (*d*_*unb*_) with 95% confidence intervals (CI) as effect size (ES) (an unbiased estimate has a sampling distribution whose mean equals the population parameter being estimated) was applied to identify pairwise differences between sexes [19]. The thresholds considered were 0.2, 0.5, and 0.8 for small, medium, and large effect sizes, respectively [21].

## Results

A total of 175 participants (52 females and 123 males) were included in the study. The mean age was 61.8 ± 6.9 years for females and 60.8±6.9 years for males, ranging from 50-76 years old.

All study participants played all the games in their friendly tournament bracket during the WF tournament. No adverse events were observed.

The descriptive and inferential results of the effect of sex on the independent variables are presented in Table 1.

**Table 1.** Descriptive and inferential analysis.

| Variables | Female | Male | Paired *t* | *p* value | Cohen *d*<sub>unbiased</sub><br>[95% CI] |
| --- | --- | --- | --- | --- | --- |
| Age (years) | 61.6±8.6 | 60.8±6.9 | -0.39 | .696 | -0.11 [-0.66; 0.44] |
| Played time (min) | 15.7±7.8 | 22.4±9.8 | 4.41 | <.001 | 0.73 [0.40; 1.06] |
| **Heart Rate** |  |  |  |  |  |
| Heart Rate Peak (BPM) | 161.5±20.7 | 161.3±17.8 | -0.06 | .955 | -0.01 [-0.39; 0.37] |
| Maximal (%HRmax) | 93.6±6.4 | 94.7±6.5 | 0.51 | .614 | 0.16 [-0.46; 0.79] |
| Average (BPM) | 140.8±20 | 139.3±18.8 | -0.37 | .714 | -0.08 [-0.49; 0.33] |
| Average (%HRmax) | 80.1±11.0 | 82.0±8.1 | 0.62 | .537 | 0.21 [-0.46; 0.88] |
| % Time in Zone 1<br>(<50%HRmax) | 2.9±8.5 | 0.1±0.6 | -1.85 | .072 | -0.62 [-1.3; 0.06] |
| % Time in Zone 2 (50-60%HRmax) | 4.5±8.6 | 1.9±4.5 | -1.31 | .198 | -0.44 [-1.12; 0.23] |
| % Time in Zone 3 (60-70%HRmax) | 11.4±21.9 | 9.1±13.8 | -0.44 | .665 | -0.15 [-0.81; 0.52] |
| % Time in Zone 4 (70-80%HRmax) | 18.3±16.5 | 23.3±18.3 | 0.83 | .413 | 0.28 [-0.39; 0.95] |
| % Time in Zone 5 (80-90%HRmax) | 42.5±28.5 | 38.8±21.7 | -0.46 | .651 | -0.15 [-0.82; 0.51] |
| % Time in Zone 6<br>(>90%HRmax) | 20.4±28.6 | 26.6±29.7 | 0.63 | .534 | 0.21 [-0.46; 0.88] |
| **Distance covered (m/min)** |  |  |  |  |  |
| Total Distance | 54.7±15.8 | 63.3±10.7 | 4.20 | <.001 | 0.69 [0.36; 1.03] |
| Distance < 4 km/h | 22.5±3.1 | 21.8±3.0 | -1.47 | .142 | -0.24 [-0.57; 0.08] |
| Distance ≥ 4 km/h | 32.6±16.7 | 41.7±12.2 | 4.03 | <.001 | 0.66 [0.33; 1.00] |

Additionally, individual differences and mean values from independent comparisons are shown in estimation plots: Figures 1, 2, and 3 for age, playing time, internal load, and distance metrics. Complementarily, Cohen’s *d*_*unbiased*_ (*d*_*unb*_) with 95% confidence intervals for all comparisons are illustrated in Figure 4.

**Figure 1.**
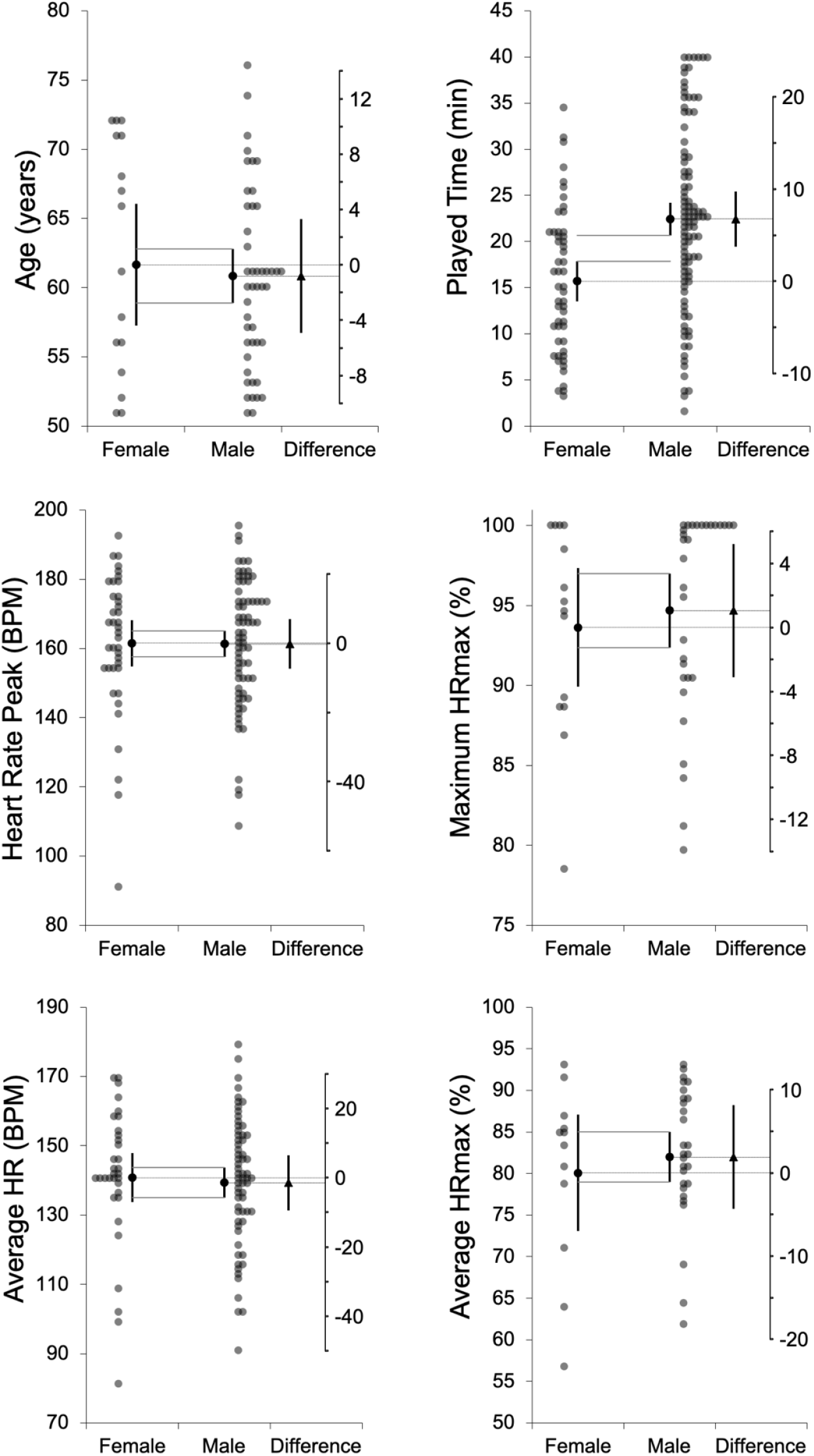
Means and 95% confidence intervals for age, playing time, and physiological variables, presented separately for females and males. The mean difference between groups, along with its 95% confidence interval, is shown on the floating difference axis on the right, aligned with the female group mean on the primary axis.

**Figure 2.**
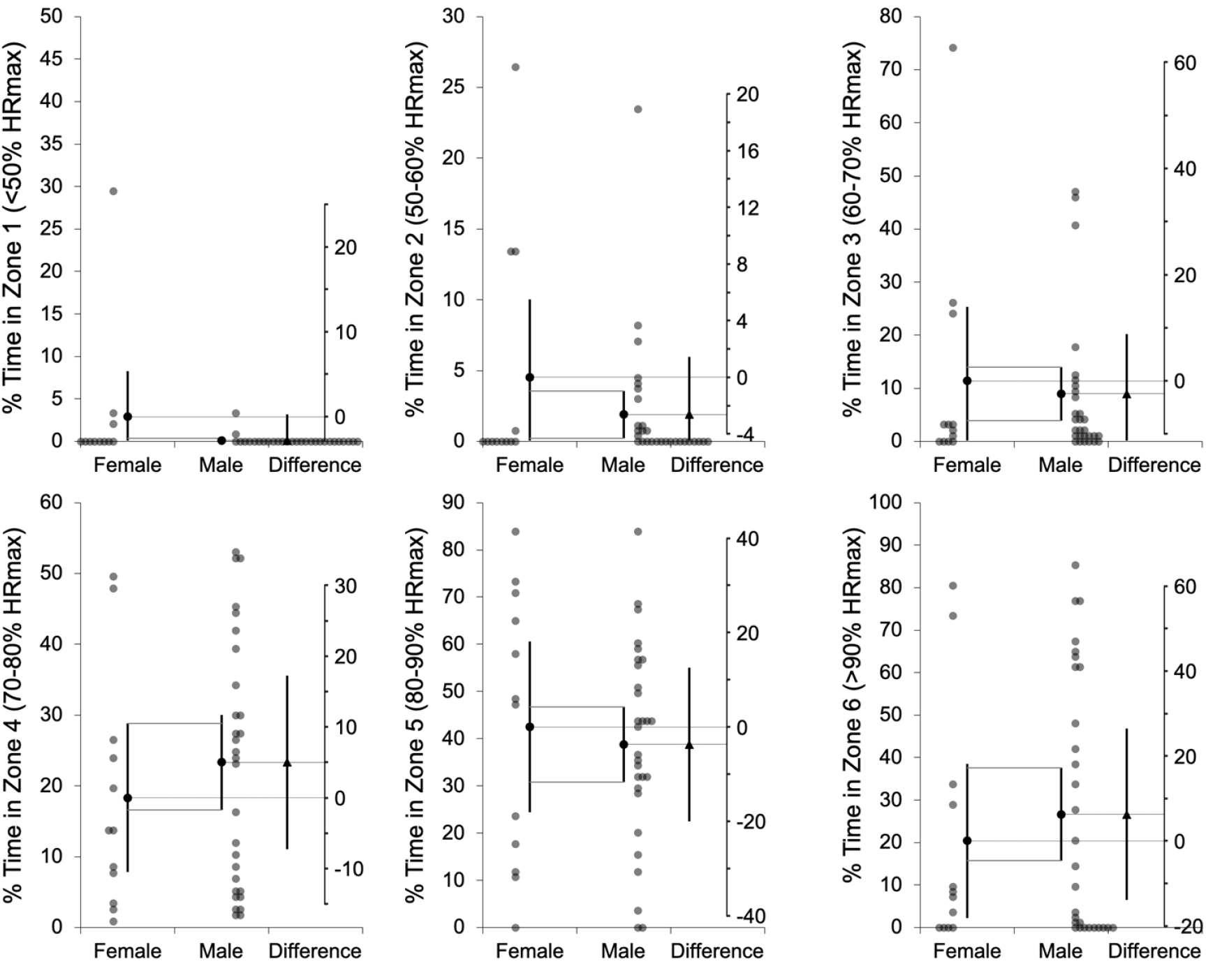
Percentage of time spent in each heart rate zone (%HRmax) across six intensity zones (<50%, 50–60%, 60–70%, 70–80%, 80–90%, and >90% HRmax), presented separately for females and males. The mean difference between groups, with its 95% confidence interval, is displayed on the floating difference axis to the right, aligned with the female group mean on the primary axis.

**Figure 3.**
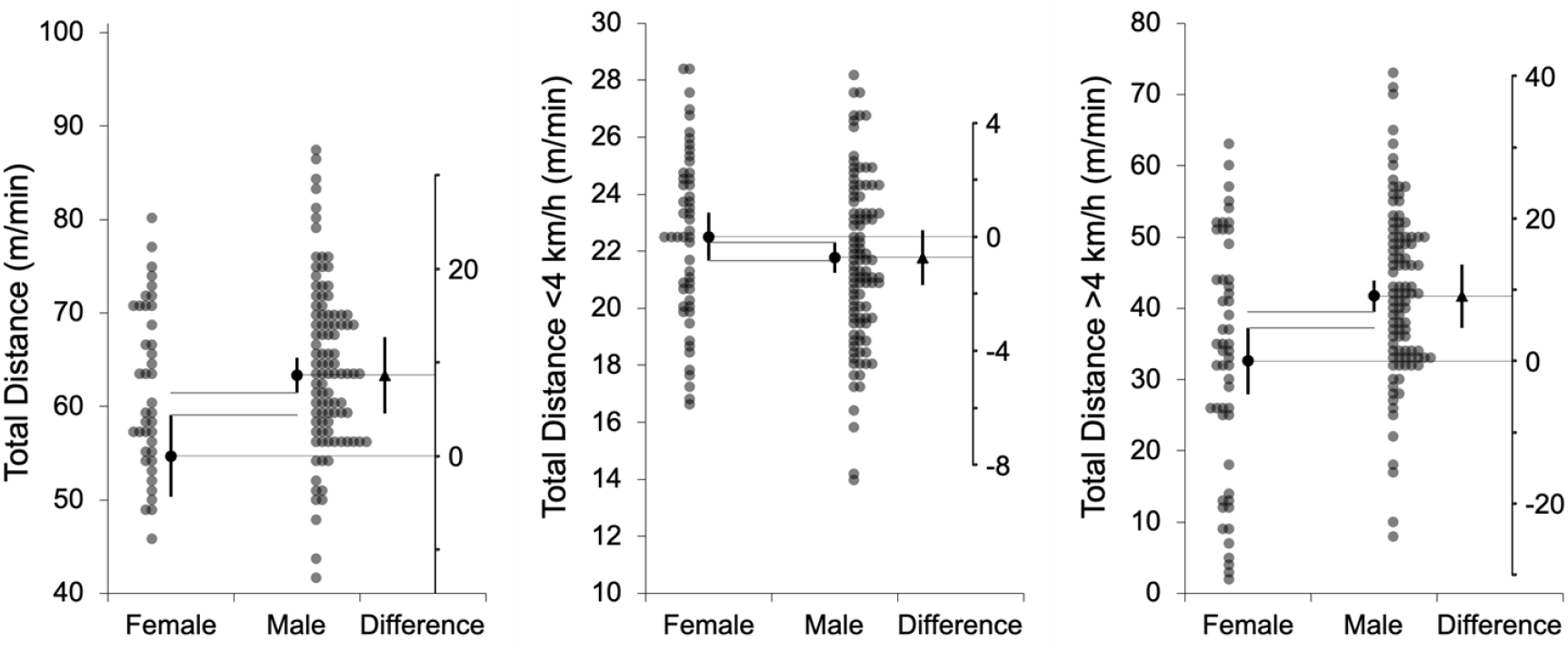
Total distance covered (m/min), and distance covered at speeds below (<4 km/h) and above (>4 km/h), presented separately for females and males. The mean difference between groups, with its 95% confidence interval, is shown on the floating difference axis to the right, aligned with the female group mean on the primary axis.

**Figure 4.**
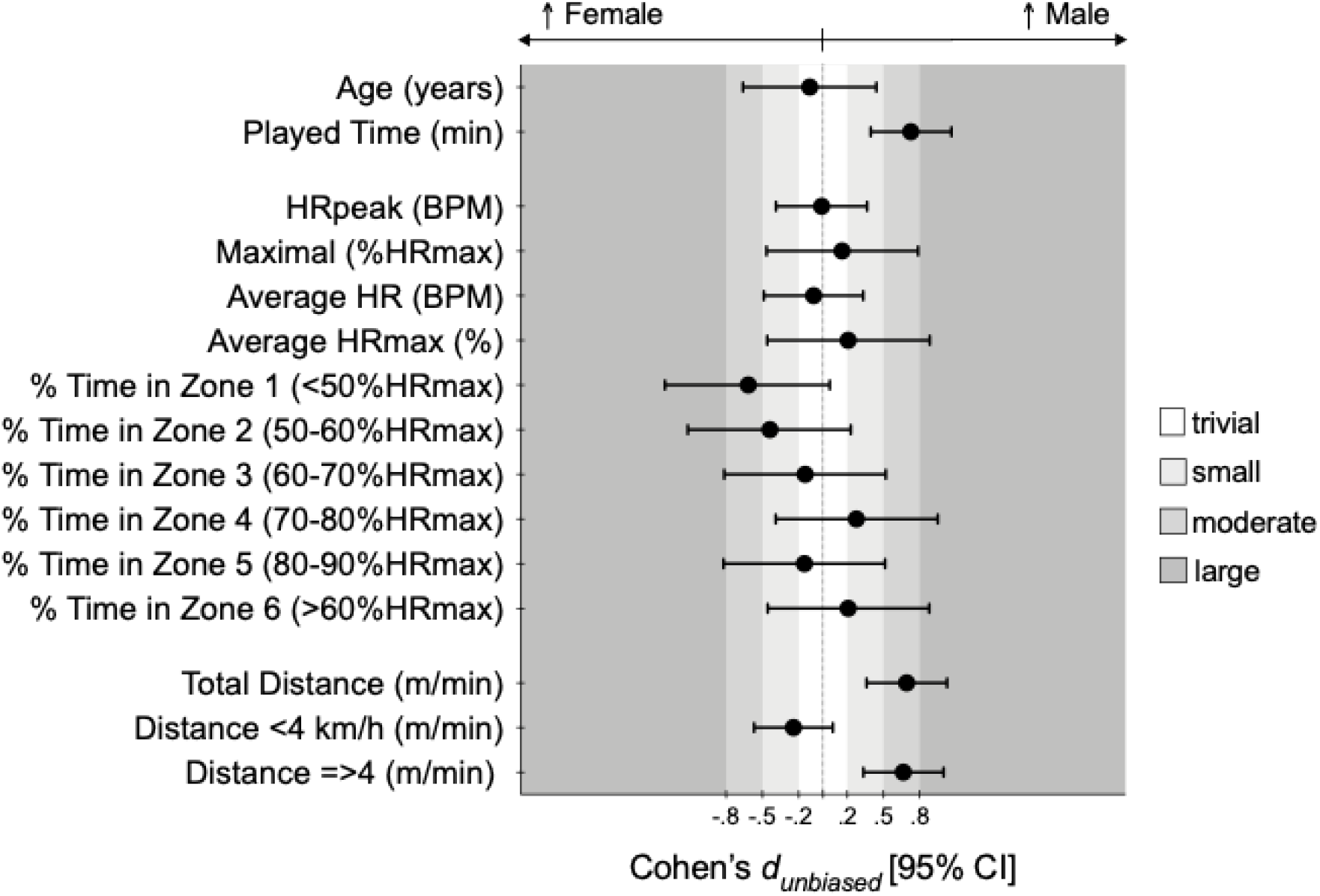
Cohen’s *d*_*unbiased*_ differences for considered according to sex. Error bars indicate uncertainty in the true mean changes with 95% confidence intervals.

Concerning external load, males covered significantly greater total distances per minute than females (female: 54.7±15.8 m/min vs. male: 63.3±10.7 m/min; t = 4.20, p < .001; *d*_*unb*_ = 0.69 [0.36; 1.03]). This difference was especially pronounced at speeds above 4 km/h (female: 32.6±16.7 m/min vs. male: 41.7±12.2 m/min; t = 4.03, p < .001; *d*_*unb*_ = 0.66 [0.33; 1.00]). Conversely, distances covered at speeds below 4 km/h did not differ significantly between sexes (t = −1.47, p = .142; *d*_*unb*_ = −0.24 [−0.57; 0.08]).

Males played significantly longer time compared to females (female: 15:41±07:46 min:ss vs. male: 22:26±09:47 min:ss; *t* = 4.41, *p* < .001; *d*_*unb*_ = 0.73 [0.40; 1.06]). Regarding internal load, no significant sex differences were observed in HR variables, including average BPM (*t* = − 0.37, *p* = .714; *d*_*unb*_ = −0.08 [−0.49; 0.33]), maximum BPM (t = −0.06, p = .955; *d*_*unb*_ = −0.01 [−0.39; 0.37]), or %HRmax across all intensity zones, such that female average %HRmax was 80.1±11.0% and male was 82.0±8.1 during the WF games.

## Discussion

The findings of this study provide valuable insights into the physiological demands of WF (i.e., analyses the external and internal load) profiles among male and female participants during a WF tournament. By examining internal and external load profiles, this research highlights the potential of WF to offer moderate-to-vigorous intensity exercise, with nuanced differences based on sex.

The results revealed that male participants exhibited higher external loads compared to their female counterparts, covering greater distances at faster speeds. This aligns with established evidence that males generally have higher absolute aerobic capacity and muscle mass, contributing to greater physical outputs during exercise [22]. However, the observed differences in external loads also underscore the need for tailored programming to ensure that WF remains accessible and beneficial for all participants. Strategies such as modifying game volume, pitch dimensions, or substitution patterns could be explored to enhance inclusivity without compromising the sport’s health benefits.

Contrary to the differences in external loads, internal loads (i.e., %HRmax) did not differ significantly between sexes. Both male and female participants reached and maintained HR within the moderate-to-vigorous intensity range, supporting the applicability of WF as a cardiovascular exercise [11, 13, 14]. These findings corroborate earlier studies that have identified WF as a suitable activity for improving heart health in older populations, regardless of sex [11, 15]. The lack of significant differences in %HRmax further reinforces the potential of WF to serve as a scalable intervention for diverse demographic groups.

WF’s ability to elicit moderate-to-vigorous intensity exercise is particularly significant given the decline in physical activity levels often observed in aging populations [8]. By meeting recommended intensity thresholds [1], [4], [5], WF can contribute to improved cardiovascular health, enhanced muscular endurance, and better weight management [14]. Moreover, this exercise modality’s structured yet flexible nature allows participants to engage at their own pace, reducing barriers to sustained participation [13].

The study’s findings also highlight WF’s potential in addressing sex-specific health concerns. For instance, females, who are more prone to osteoporosis and balance-related issues, may benefit from the sport’s low-impact movements that promote bone density and stability [23]. Conversely, males, who often face greater risks of cardiovascular events can influence WF’s aerobic benefits to mitigate such risks. Moreover, WF had positive health effects on BMI and blood pressure [24]. Furthermore, it was found that WF programmes attained moderate-intensity exercise and several changes of direction which can produce benefits on cardiovascular and metabolic health and osteogenic stimulus [24].

While this study offers robust insights, certain limitations warrant consideration. The cross-sectional design excludes causal inferences about the long-term health impacts of WF. Additionally, although the sample reflects the proportion of men and women participating in the WF tournament from all districts of Portugal, the number of women was significantly lower, which may also limit the generalizability of the findings.

It is worth highlighting as a strength that this study is the first to evaluate measures associated with internal and external load in a WF tournament context. The study successfully provides an analytical characterization of these variables for both male and female undergoing the aging process, considering the international physical activity guidelines. Future research should adopt longitudinal designs to explore the sustained effects of WF on health outcomes and incorporate diverse populations to examine cultural and environmental influences on participation. Moreover, exploring individualized adaptations to WF, such as individual intensity levels/thresholds or modified game rules, could enhance its efficacy and safety for participants with varying fitness levels and health conditions.

## Conclusion

Overall, although males generally played more time than females and covered a higher distance per minute, showing a higher external load, both males and females report similar internal loads ranging from moderate to vigorous exercise intensity. By addressing sex-specific demands and promoting health, WF holds significant promise as a scalable intervention for aging populations. Continued research and innovation in WF programming are essential to fully harness its potential and extend its reach to broader populations.

## Acknowledgments

The authors would like to thank the participants for their participation and cooperation during the study.

## Author contributions

Conceptualization, Methodology, Formal Analyses, Writing - Original Draft: Júlio A. Costa Catarina Pereira, André Seabra and Bruno Gonçalves; Methodology, Writing - Review & Editing: Júlio A. Costa, Catarina Pereira, André Seabra, Ana Barbosa, Ana Pinto, Catarina Martins, Rafaela Moreira and Bruno Gonçalves; Supervision, Conceptualization, Methodology, Writing - Review & Editing: Júlio A. Costa and Bruno Gonçalves.

## Data availability statement

Data cannot be shared publicly because the data contain potentially sensitive information and data have been obtained from a third party (i.e. Portugal Football School, Portuguese Football Federation) and access restrictions apply. Data are available from the Data Protection Office, Portuguese Football Federation (Data Access contact via) for researchers who meet the criteria for access to confidential data. The authors confirm that they did not have any special access privileges that others would not have and all relevant data can be accessed in the same manner as the authors, from the Data Protection Office, Portuguese Football Federation Data Access.

## Code availability statement

Not applicable.

## Ethics approval and informed consent

All participants provided written informed consent prior to participation, and the study was approved by the Portugal Football School, Portuguese Football Federation (Protocol number: 21/CEPFS/2023).

## Declaration of Generative AI and AI-assisted technologies in the writing process

During the preparation of this work the authors used the Open ChatGPT to compose small parts of the text and to translate it into English language. After using this tool, the authors reviewed and edited the content as needed and take full responsibility for the content of the publication.

